# Expansive growths with uniaxial gradients can explain formation of oblong diversity observed in two-dimensional leaf shapes

**DOI:** 10.1101/2022.08.06.503069

**Authors:** Akiko M. Nakamasu

## Abstract

Regulation of positional information in fields with different sizes are known as scaling in the area of morphogenesis, and enable integrated and robust developmental processes. Although it is known that interpretation of such scaled patterns leads to formations of relative shapes, the same positional information brings about diversities in morphogenesis.

In this research, a boundary of a two-dimensional shape was constructed by propagating points and segments connecting them for a description of a growing form. Cell expansion “with” or “without” cell proliferation were implemented using different simple algorithms, as “additive growth” and “expansive growth”, respectively. When the different types of growth algorithm with a biased restriction were calculated, the additive growth maintained a relative shape corresponding to the gradients with different lengths. However, diverse shapes were generated by the gradients in the cases of expansive growth and its combinations even with negate effects by additive growth. As an operative example of this attempt, leaf shapes with smooth margins were calculated using a combination of these growth algorithms.

Finally, we concluded that different algorithms brought different responses against the simple positional information, i.e., additive growth always governed by it or expansive growth can escape it. It was predicted theoretically that an expansive growth has a capacity to become a generator of diversity at least in leaf morphogenesis.

**Summary statement:** Explanation of different responses against single positional information by different algorithms for two types of growth modes.

## Introduction

Developmental processes are reproduced. For this purpose, a positional information need to be read out identically. It is well known that small plteus larvae can be obtained from half embryos of sea urchin (Driesh, 1892). Similar phenomena have also been observed in Salamandridae embryos (Spemann, 1938), among other organisms. Therefore, robust shaping can be achieved even with different sizes of positional information. If so, how we can obtaine diversity?

In the context of such regulative developments, dynamic adjustments of axial patterns to embryonic sizes have been reported (Cooke, 1981). Such regulation of positional information is known as “scaling” (Schmid-Nielsen, 1984). The mechanisms for the scaling of positional information have been treated theoretically (Murray, 2001), then investigated in several model systems on a molecular level as summarized in (Capek and Müller, 2019).

To obtain robust shape from such scaled patterns as positional information, relative shaping is needed during outline-shaping processes. Cell or tissue determination or differentiation processes based on positional information were also discussed theoretically as interpretation of French Flag by (Wolpart, 1969), and a biochemical analogue-to-digital converter by (Meinhardt, 1982), among others. Experimental systems, e.g., a proximo-distal sequence of cartilage structure in chick limbs (Summerbell, 1973; Morishita *et al*., 2014), and insect legs, and so on were selected to address this problem. However, information regarding the robustness in two-dimensional outline-shaping processes is still insufficient.

Formations of plant leaves also follow the shaping robustness. However, at a same time, morphological diversities are derived from differences in positional information. For example, it is known that a difference in the boundaries of basal growth zones influence on a leaf-shape complexity through a control of marginal outgrowths in their formation processes (Kierzkowski *et al*., 2019). Then leaf-shape diversities can also be observed in simple leaves with an entire margin. For example, the diversity of foliage leaves even within a single plant is known as a heteroblasty in *Arabidopsis thaliana* (Tsukaya, 2000). Then the WOX related change in proportion was suggested in (Zhang *et al*., 2020). Such diversities may also be obtained by similar differences in positional information. In this current work, we explore mechanisms by such proximodistal positional information might affect leaf morphogenesis.

In this research, a simple two-dimensional system of shape boundary was utilized for descriptions of growing forms, because of the complexity in three-dimensional morphogenesis including many information and events. The modeling of a leaf formation was tried in Kuchen *et al*., 2012 as mentioned cell growth on blade other than peripheral, and importance of feedback effect was suggested in Hervieux *et al*., 2016. The description of a shape boundary for simple leaf with serration was done in (Bilsborough *et al*., 2011). Then a similar boundary method constructed by propagating points and segments connecting adjacent points was previously proposed in (Nakamasu *et al*., 2014).

In plant morphogenesis, that lack almost cell movements, different growth modes caused by cell expansions with or without cell proliferations mainly contribute to the resulting shapes. Therefore, effects of these biological events on a shape formation were expressed by simple algorithms as additive growth and expansive growth. A linear gradient starting from the base of a shape was set as the simplest positional information. Then the effects of difference in gradient lengths were investigated about the respective growth algorithms. Finally, as an operative example of a combination of these growth modes, leaf formations without the marginal indentations were calculated.

## Materials and methods

### Algorithms for additive and expansive growth

Follow a boundary propagation method (Sethian 1999) that is, space propagation over time for geometrical deformations, we utilize boundary description in this paper. In our method, a contour is expressed discretely (i.e., by segments and connection points of them) then propagation is applied iteratively by updating the connection points (Nakamasu *et al*., 2014). When the lengths of the segments exceed a threshold through the propagation, the segments are divided into two. The connection point 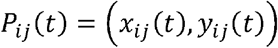 of the adjacent *i*th and *j*th segments is displaced as 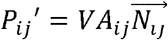. Here, for a description of the effect by cell proliferation, *VA_ij_*(*t*) is a velocity and 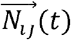 is a local unit vector on an apex pointing outward from a closed contour. (Fig. 1A). This rule for boundary movement is fundamentally different from Bilthborough et al. 2011. The cell proliferation results in a local deformation, subsequently brings a hebetate protrusion. Such a shape can be observed in leaflets of *Eschscholzia californica* primordia with similar sizes of cells on the tips (i.e., they seem to be caused by cell proliferation) (Ikeuchi *et al*., 2013). The value of *VA_ij_*(*t*) is assumed to be affected by morphogen concentrations at that time, for example, *u_i_* and *u_j_* in adjacent segments on the contour and/or *w_xy_* that is determined in a position unrestricted on the contour. In this research, it only depends on *y*, i.e., *w_y_*. Therefore, the growth speed at the point *P_ij_* is determined as a function of (*u_i_* + *u_j_*) and/or *w_y_*. For description of the growth caused by cell expansion without cell proliferation had been treated in (Nakamasu *et al*., 2017), though, for its biased case, the connection point *P_ij_*(*t*) is propagated along the vector from the geometrical center of the initial condition as 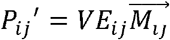. Here, 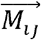 is defined as the vector on each point in the direction mentioned above and with a length of the distance from the origin is divided by a representative length *l* of the shape (*l* is the length of the tallest vector) (Fig. 3A). Then *VE_ij_*(*t*) is also affected by *w_y_* in this research. Calculations are done on the platform Wolfram Mathematica ver. 12.1.1. Then parameters αs, βs, and γs utilized in simulations are shown in Table 1. Resulted shapes without biase are obtained in every 5000 iterations then superimposed without alignment in Fig. 1B, J, Fig. 3B.

**Fig. 1.**
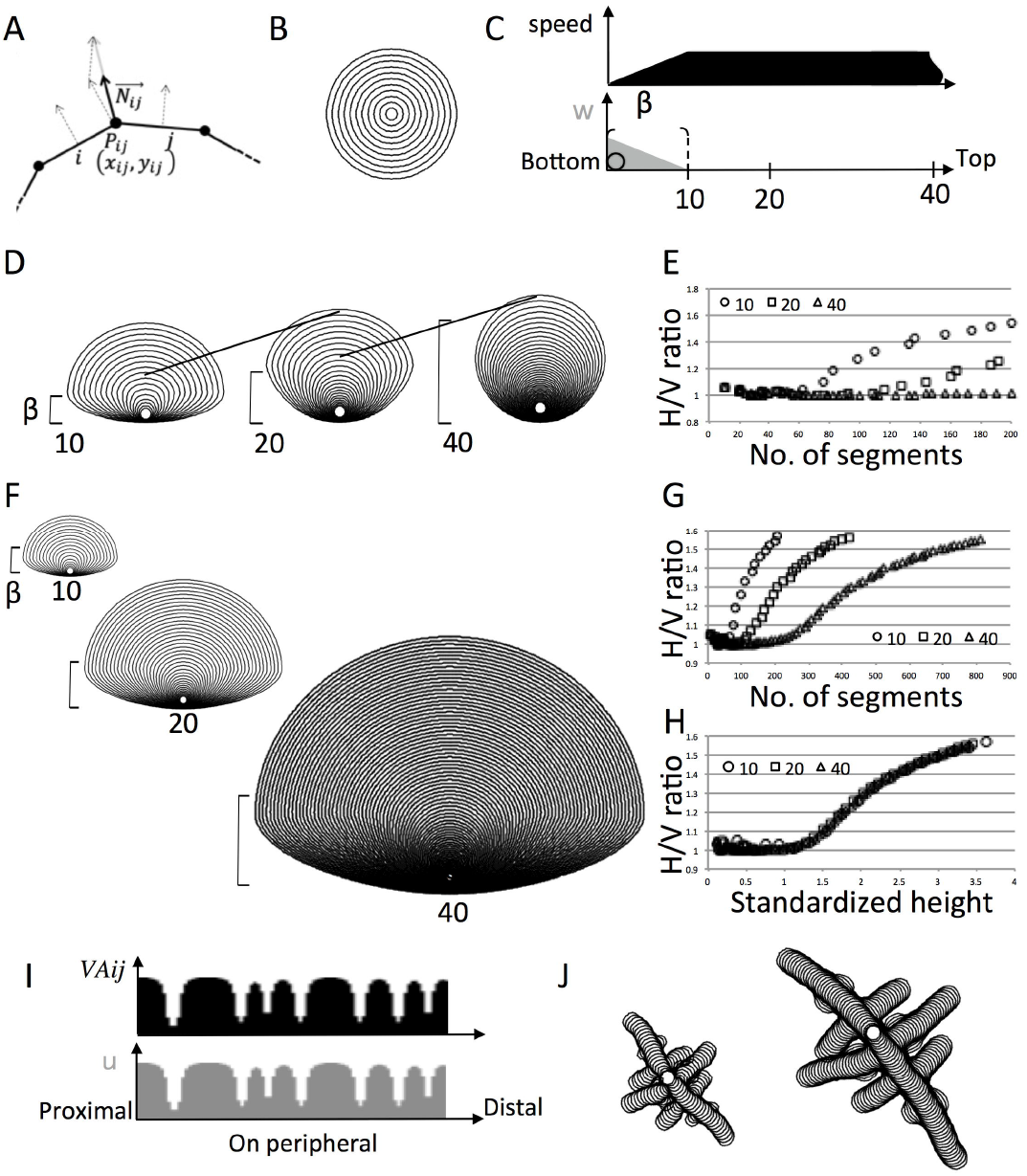
An algorithm for expansive growth. (A) Schematics of propagation of a connection point. Deformation rules of a polygon in additive growth algorithm was shown. A connection point of two segments that composes a contour was moved along a unit normal vector of the apex (B) Superimposed time series of shapes obtained by the additive growth. (C) Schematics of the given gradient and its function on growth speed are illustrated. The concentration of the substance *w_y_* linearly decreased along the vertical axis from the base of the shape. It functions as a biased restriction of growth. Then the size of initial condition was superimposed in it. (D) Superimposed time series of shapes obtained by a geometrical sequence of β = 10,20,40 in the additive growths. Different lengths of the inhibitory gradients resulted in a nested similarity. (E) H/V ratios plotted against number of segments. (F-H) Shaping robustness corresponding to the lengths of inhibitory gradient with different length β = 10,20, and 40. (F) Superimposed time series till segments had reached the values of 2^*n*–1^ of 200 segments for respective *n* = 1,2,3. Horizontal/vertical ratios were plotted against (G) the number of segments in each contour and (H) standardized heights of the shapes. (I, J) Shaping robustness corresponding to the wave lengths of periodicity that inhibit the additive growths. (I) Schematics of prophyles of *u*-distribution and *VA_ij_* on the peripheral. (J) Superimposed time series of boundaries obtained by an additive growth with periodical inhibition on the margin. The periodic patterns as each positional inhibition were generated by a reaction-diffusion system with different scale parameters (Murray, 2001).

**Table 1.**
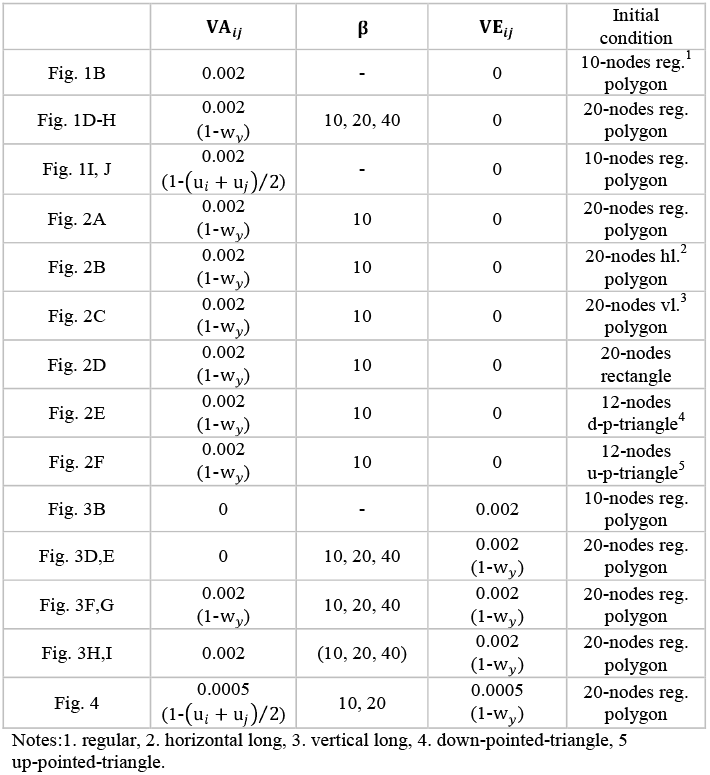
Parameters utilized in the biased restriction of growth.

| | $VA_{ij}$ | $\beta$ | $VE_{ij}$ | Initial condition |
| --- | --- | --- | --- | --- |
| Fig. 1B | 0.002 | - | 0 | 10-nodes reg.<sup>1</sup> polygon |
| Fig. 1D-H | 0.002<br>(1-$w_y$) | 10, 20, 40 | 0 | 20-nodes reg. polygon |
| Fig. 1I, J | 0.002<br>(1-($u_i + u_j$)/2) | - | 0 | 10-nodes reg. polygon |
| Fig. 2A | 0.002<br>(1-$w_y$) | 10 | 0 | 20-nodes reg. polygon |
| Fig. 2B | 0.002<br>(1-$w_y$) | 10 | 0 | 20-nodes hl.<sup>2</sup> polygon |
| Fig. 2C | 0.002<br>(1-$w_y$) | 10 | 0 | 20-nodes vl.<sup>3</sup> polygon |
| Fig. 2D | 0.002<br>(1-$w_y$) | 10 | 0 | 20-nodes rectangle |
| Fig. 2E | 0.002<br>(1-$w_y$) | 10 | 0 | 12-nodes d-p-triangle<sup>4</sup> |
| Fig. 2F | 0.002<br>(1-$w_y$) | 10 | 0 | 12-nodes u-p-triangle<sup>5</sup> |
| Fig. 3B | 0 | - | 0.002 | 10-nodes reg. polygon |
| Fig. 3D,E | 0 | 10, 20, 40 | 0.002<br>(1-$w_y$) | 20-nodes reg. polygon |
| Fig. 3F,G | 0.002<br>(1-$w_y$) | 10, 20, 40 | 0.002<br>(1-$w_y$) | 20-nodes reg. polygon |
| Fig. 3H,I | 0.002 | (10, 20, 40) | 0.002<br>(1-$w_y$) | 20-nodes reg. polygon |
| Fig. 4 | 0.0005<br>(1-($u_i + u_j$)/2) | 10, 20 | 0.0005<br>(1-$w_y$) | 20-nodes reg. polygon |

Notes:1. regular, 2. horizontal long, 3. vertical long, 4. down-pointed-triangle, 5 up-pointed-triangle.

### Analyses for biased growths

For a biased inhibition of the growth speed, *w_y_* decreases linearly corresponding to the *y*-coordinate from the base *y_b_*(*t*), and eventually drops to zero at the distal part of the shape, as shown in Fig. 1C,

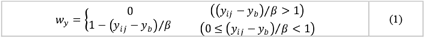

Then, 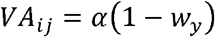 for an additive growth and *VE_ij_* = *γ*(1 – *w_y_*) for expansive growth are utilized in this research. Both modes are constantly repressed by the *y* dependent linear gradient *w_y_*, in the same form, but via different constants α and γ, respectively. As an initial condition, a regular polygon composed of 10, or 20 nodes is utilized. Resulted shapes are obtained in every 5000 iterations then superimposed aligning the bottoms in Fig. 1D, F, and Fig. 2A-C, E-G, and Fig. 3D, F, H.

**Fig. 2.**
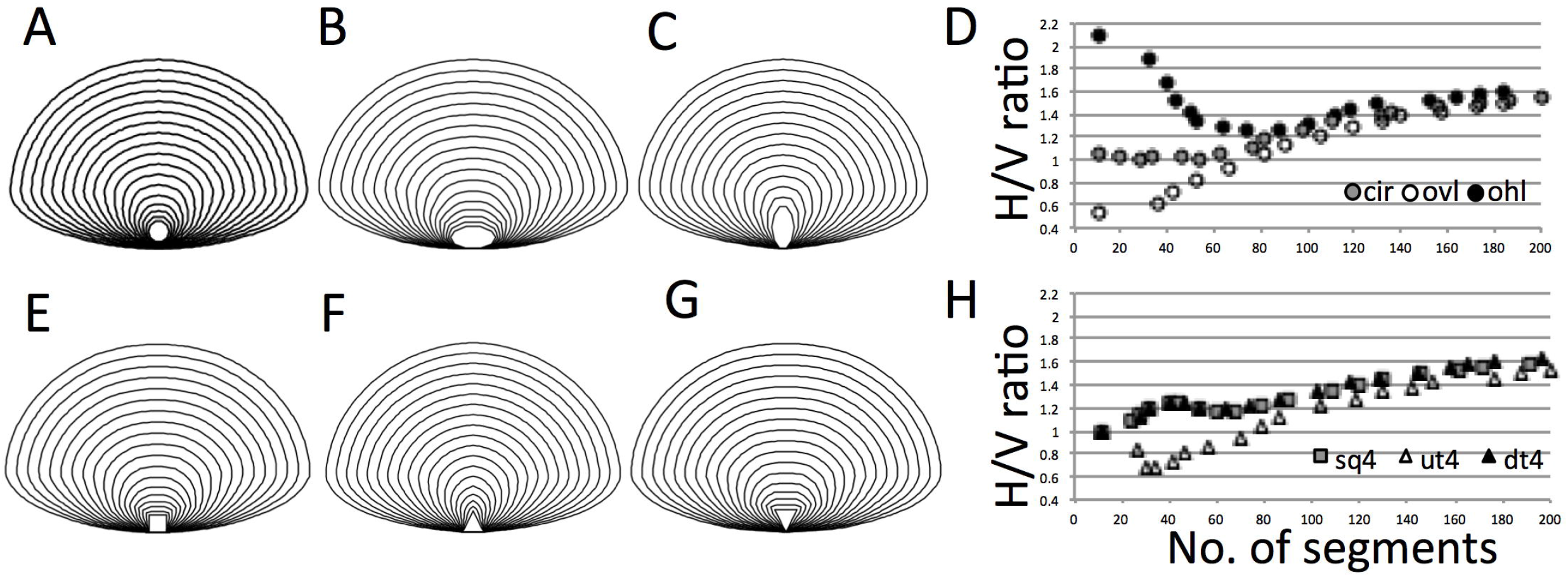
A shaping robustness was observed against a shape difference in initial conditions. Examples of superimposed time series of shape boundaries from each trial of additive growths. Simulations were started from initial conditions with different shapes as (A) regular polygon, (B) wide- and (C) tall-polygons. (E) square, (F) up-ward-pointed triangle, and (G) down-ward-pointed triangle. (D), (H) Horizontal/vertical ratios plotted against the number of segments in each contour.

**Fig. 3.**
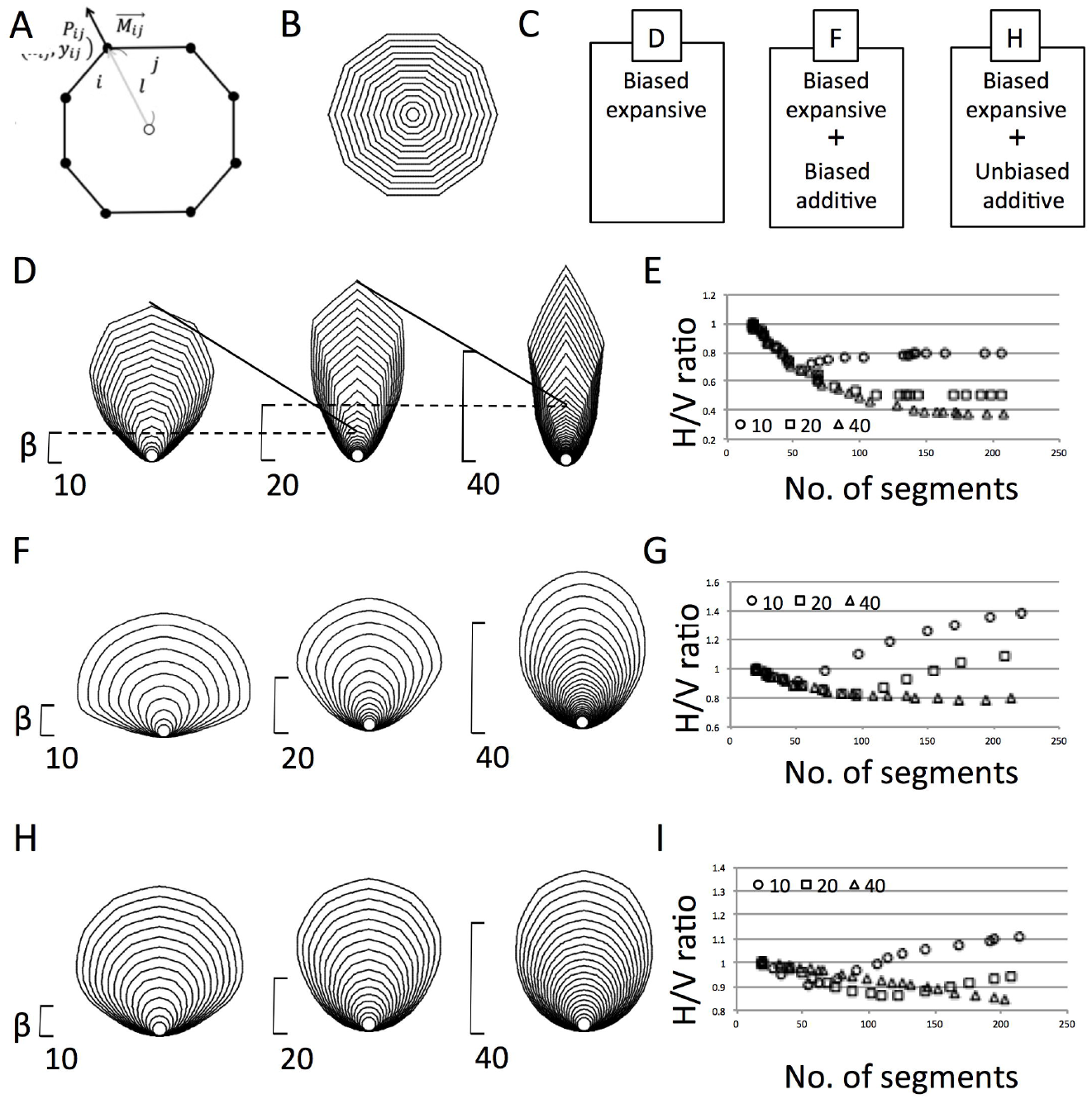
An algorithm for expansive growth and combination with additive growth. (A) Schematics of propagation of a connection point. Vector from the geometrical center of the initial condition in expansive growth algorithm. (C) Schematics of the biased expansive growth and their combinations. (D, F, H) Superimposed time series of shapes obtained by a geometrical sequence of β = 10,20,40 in the expansive growths (D) and combinations with additive growths with biase (F) and without biase (H). (E, G, I) H/V ratios plotted against number of segments. (E) expansive growth, and combinations with biased (G) or unbiased (I) additive growths.

### Simulation for leaf-like shapes

Following the methods of Harrison and Kolar in 1988, a Turing pattern of a reaction-diffusion (RD) system (Turing, 1952) is utilized to implement an arbitral periodicity, although the pattern is thought to be biologically induced by a polar auxin transport (PAT). The utilized condition have a critical wavelength at a selected parameter set, and the wavelength is stable against growth. The partial differential equations used in the simulation have linear type reaction terms with limits (Kondo and Asai, 1995).

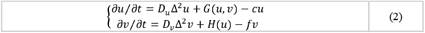

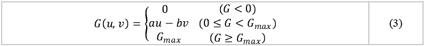

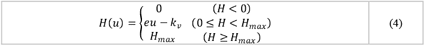

The parameters used in the simulation are as follows; *dt* = 0.2, *ds* = 1, *D_u_* = 0.015, *D_v_* = 0.45, *a* = 0.04, *b* = 0.04, *c* = 0.012, *e* = 0.07, *f* = 0.05, *k_v_* = 0.025, *G_max_* = 0.01, and *H_max_* = 0.05. The periodical positional information by RD is espressed by different concentrations of component *u* on respective connected segments *i*th *j*th, i.e., *u_i_*, *u_j_*, …etc. Then the pattern is affected by the gradient *w_y_* that work on the diffusivity *D_u_* as *D_u_*/(0.1 + 0.9*w_y_*). Therefore, the wavelength become long and finally pattern disappear for the deficient of *w_y_*. This would strongly change the local outgrowths of the contour. The parameter-set yields a stable splitting with domain extensions as similar shown in (Nakamasu *et al*. 2014).

For a leaf-like shape, a combination of above two growth algorithms is utilized. As mentioned in (Bilsborough *et al*. 2014), the propagation is periodically changed on the boundary, in the model for leaf serration of *A. Thaliana*. In this research, the boundary propagation of additive growth is periodically inhibited by the periodic pattern. A gradient, that composed in the leaf model by (Bilsborough *et al*. 2014) inhibited a periodical growth on the margin, although, the gradient activates the periodical pattern formation in this research (i.e., activation of the biased restriction of additive growth with periodicity), and the gradient also inhibits the expansive growth. The speeds set as *VA_ij_* = *α*(1 – (*u_i_* – *u_j_*)/2) for periodical additive-growth, then *VE_ij_* is same as the form mentioned above for the expansive growth. Resulted shapes are obtained in every 10000 iterations then superimposed aligning the bottoms in Fig. 4A, B.

**Fig. 4.**
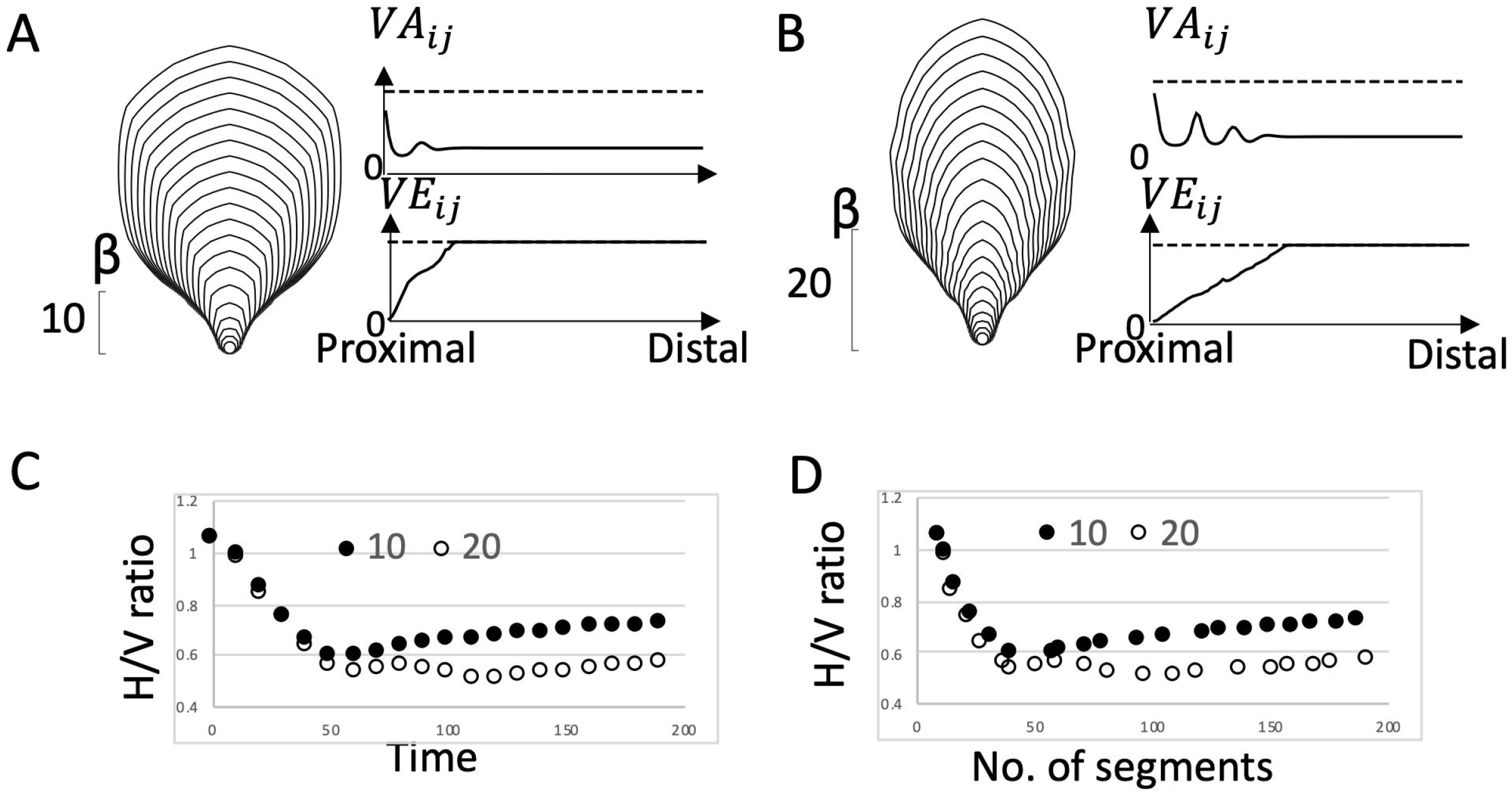
Leaf-like shapes with different proportions generated by inhibitory biases in combination of additive- and expansive-growths. Examples of superimposed time series of shape boundaries obtained by a simulation result of leaf like shape (A, B). (A) Shorter bias with β = 10 and (B) longer bias with β = 20. The gradient lengths were set within a triple the length of marginal periodicity. In their right-hands, distributions of two kinds of speed (*VA* and *VE*) in each last frame were shown in respective graphs. These distributions along the margin from the proximal to the distal of each shape were plotted. Uppers are *VA_ij_* for additive growth and lowers are *VE_ij_* for expansive growth. Dashed lines indicate *α* of *VA_ij_* and γ of *VE_ij_*, respectively. (C) The time-series plot of H/V ratios and (D) H/V ratios plotted to the number of segments in the contour.

## Results

Plant morphogenesis is known to lack drastic cell movements. Therefore, in cell based growth the existence of cell proliferation greatly affect the shaping processes. Boundary growth keeps the polygon with straight line by expansion, though cell proliferation can absorb the straightness with local deformation. The difference in the involvement of cell proliferation was implemented in a boundary of a growing form by simple algorithms as different growth modes (Fig. 1A, Fig. 3A). The algorithms are named as additive growth and expansive growth, respectively. The shapes that can be generated by these algorithms were explored, as follow.

### Relative shapes were obtained against uniaxial positional information with different lengths in the additive growth algorithm

In the algorithm for additive growth, a contour was expressed by the closed polygon composed of a set of points and segments, what connect two adjacent points. Each point propagated over time to the centrifugal side along its normal vector (Fig. 1A). When the length of a segment exceeded a threshold by the propagation, the segment was divided. If an initial condition was given as a regular-polygon arrangement composed of 10 nodes, the shape grew with a constant *α* as the propagation speed. The propagated boundaries keeping smooth curves were superimposed in Fig. 1B as a time-series.

Next, a uniaxial linear-gradient with a length *β* was added as stable positional information to this system (Fig. 1C). Subsequently, a symmetrical growth from a regular polygon (includes 20 nodes) was biased (Fig. 1D). If the gradient (starting from the base) had a function to suppress the growth, the shape grew laterally. It was caused by a continuous suppression of the local growth within the *β* height. As a result, it became an oval shape with a long horizontal-axis. Three trials of calculation against a geometric sequence of *β_n_* = *βr*^*n*–1^ (*β* = 10, *r* = 2, *n* ≤ 3) were shown in Fig. 1D. The calculations were continued until the number of segments over 200, in these trials. The time-series with their respective *n*’s seemed to include similar shapes. A contour with a different size was observed in an earlier time of another time-series with a smaller *n* (i.e., these time-series till a certain time seemed to be nested). When the calculation of each trial was extended until the number of segments had reached the values of *r*^*n*–1^ of 200 for respective ns, the boundaries had an asymptotically equivalent shape (Fig. 1F).

When horizontal to vertical (H/V) ratios were obtained as an allometric index of these shapes, the H/ V ratios increased with an increase in the number of line segments (Fig. 1E, Fig. 1G). Then, the overlap of the plots could be confirmed when the lengths were standardized with *β* (Fig. 1H). Therefore, obtained shapes were relative to the *β*s; each length of gradient.

In the case of a periodical growth on the boundary (Fig. 1I), the relative shaping can also be obtained (Fig. 1J). A characteristic rule mentioned in (Nakamasu *et al*., 2014; Nakamasu *et al*., 2017, Nakamasu and Higaki, 2019) was followed in the branch arrangements caused by positional information with different wave lengths of periodicity.

Therefore, this additive growth algorithm seemed to give similar shapes in the morphogenesis. That is, relative shaping could be observed as the percentage of *β* that was determined by *w_y_*-related positional information.

### A shaping robustness was observed against different shapes of initial condition in the additive growth algorithm

Although the shape was updated at each step, relative shapes could be obtained to the length of gradients for the uniaxial restriction of additive growths. These growing shapes were regarded as robust against difference in sizes of initial conditions, and time- and spatial-intervals for calculations, i.e., the propagation speed α and the segment threshold *th* with *β*-scalings, in a certain range.

Considering about initial conditions, symmetrical growths from a regular polygon with 20 nodes weren’t hardly disturbed within the early calculation steps. Because the effects of the gradient were small enough on a small shape when it was compared with the gradient. Even in the cases of the polygon shape as an initial condition was changed to wider or taller polygonal ring, shaping robustness was also observed (Fig. 2A-C). However, obtained shapes were slightly affected by the initial conditions (Fig. 2D). A 20-nodes polygon with a 2:1 H/V ratio resulted in a longer horizontal axis than the others, and a polygon with a 1:2 H/V ratio resulted in a shape with a longer vertical axis than the others. When the simulation was started from a square or an up- or a down-ward-pointed triangles, the obtained shape also affected by the initial condition (Fig. 2E-G). Though, they approached the asymptotically equivalent shape that starting from a regular polygon (Fig. 2A). That is, the robustness during shaping processes could be obtained against different shapes of initial conditions.

A comparison of plots of aspect ratios in these shapes showed a greater overlap between the down-ward-pointed-triangle case and the square case (Fig. 2H). It was obvious that the lower side (with less propagation by a biased restriction) was more robust to the change in shapes.

Therefore, it was found that a uniaxial gradient was sufficient to regulate a two-dimensional shape. It was then considered that the restrictions of growth speed were effective for the shaping robustness.

### A shaping variation was obtained by a difference in lengths of inhibitory gradients in this expansive growth algorithm

In the case of expansive growth, connection points in a closed polygon were propagated along the vectors from the geometric center of the initial condition. As a result, divided triangles in the polygon that share each apex at the geometrical center were magnified then the growth kept shape similarity. A set of 10 segments arranged as a regular polygon for the initial condition, the shape was simply magnified, although the edges composed of some line segments because, initial segments were divided in several times into two segments when it exceeded a threshold. They sustained straightness (Fig. 3B), while additive growth bend it. Even though each propagation proceeded at a constant rate *γ*, the magnification process was accelerated by extending vectors referenced. To remove the acceleration, the vectors were normalized by the representative; i.e., tallest, vector.

The time-series of growing shapes by the expansive growth with inhibitory gradients were shown in Fig. 3D. A uniaxial linear-gradient with a length *β* was added as positional information to this system. Three trials of calculation for each *β* in geometric sequence of *β_n_* were done. Each initial polygon (composed of 20 nodes) seemed to grow taller with the biased growth. Because the differences between the lengths of adjacent vectors were emphasized during the growing shape within the gradient.

When the aspect ratio was plotted against the number of segments in a contour, the time series was decreased from a value of 1 for extension along the vertical axis. The declining plot showed a similar transition rate within each gradient (Fig. 3E). Subsequently, when the growing shapes escape the gradient, they started to expand maintaining the proportion at that point. Therefore, boundaries with a similar shape were commonly observed among these time-series with the respective *n*’s. A shape with a different size observed at an earlier time of the time series with a larger *n* (i.e., the time series seemed to be nested in reverse order of the case of additive growth).

Therefore, in this expansive algorithm, it was found that the difference in the lengths of inhibitory gradient will be converted to a diversity in shaping.

### The shaping variation caused by the expansive growth algorithm was not cancelled even in combinations of additive growths

When the expansive growth was combined with additive growth with or without biase (Fig. 3C), the time-series of shapes obtained by different *β_n_*, showed no longer similar (Fig. 3F-I). The shape variation, i.e., oblong diversity caused by the expansive growth algorithm, were not cancelled even with the negate effects by the additive growths. Therefore, the changing shape did not show a shaping robustness corresponding to the length of the gradient. In the H/V ratios plotted against the number of line segments within a contour, the ratios first decreased with overlap then increased at respective timings, as shown in Fig. 3G, I. A larger *n* thus has a longer vertical axis within these ranges.

In the case ofleaf shape formation, such a combination of growth modes seems to be operative. Mixtures of cell sizes observed in marginal serrations in *A. thaliana* leaves (Kawamura *et al*., 2010) indicate the existence of cell expansions accompanied by cell proliferations i.e., a combined growth modes mentioned above. As shown in (Bilthborough *et al*., 2011), periodical growth at the margin and the bias of the growth rate were incorporated together for description of leaf development. Difference in the biased positional information is known to be important for a particular shaping in actual leaves. That is, the boundary of the graded basal growth zone can change the complexities of the leaf shape (Kierzkowski *et al*., 2019).

It was considered the cases even in entire leaves. When the gradient lengths were fixed within about double wavelengths (i.e., *β* ≲ 2*λ*) of the periodic pattern, entire leaf-like shapes with smooth margin could be obtained autonomously. *λ* is the critical wavelength of a Turing pattern that caused by interactions between *u* and *v* (Miura and Maini, 2004), then it was derived as ~8.8 in this parameter set. In Fig. 4A, B, both leaf shapes were simple within the entire margins, although, these leaves indicated different proportions by different *β*s. The longer bias resulted in a narrower proportion, as expected from the above result of combination in Fig. 3F, H. The change in the aspect ratios (Fig. 4C, D) were also follow the result of combined case in Fig. 3G, I. These effects of additive growth were not linear in this case (as shown in the graphs in Fig. 4A, B), though, a difference in the proportions was obtained certainly by the expansive growth that encouraged by inhibitory gradients with different lengths.

It was confirmed that when the expansive growth read out inhibitory gradients, it became a generator of diversity in terms of shape, as well as in the case of the combination of the algorithms. Leaf-like shapes with different proportions were regenerated by different lengths of the gradient in simulations of combined growth modes.

## Discussion

In this research, different algorithms, based on different growth modes, and their combinations were examined to considering the morphogenetic problems related to response to positional information. These growth modes, cell expansion with or without cell proliferation, were implemented with simple algorithms, additive and expansive, respectively. During these trials, uniaxial gradients with different lengths were given as the positional information for biased restrictions of additive (Fig. 1) and/or expansive (Fig. 3) growths.

In the additive growths, relative shapes correspond to the length of the gradient was observed (Fig. 1D-J, Fig. 2). The relative shaping was maintained against spatial- and temporal-difference in intervals of the calculations (Fig. 1F-H). Then the shaping similarity could also be obtained against different lengths of periodicity on the margins (Fig. 1I,J). It may seem geometrically self-evident, though the robustness of growing shapes from different sizes and shapes of initial conditions (Fig. 2) indicates a two-dimensional effect on shaping by positional information of uniaxial gradient. The obtained shape resembled a bacterial colony grown on inhomogeneous environment (Tasaki *et al*., 2017), that seems to be brought by the proliferations of linked cells on the edge. Such regularities to positional information may follow the morphogenetic robustness in development and regeneration as described in (Thompson, 1917; Niklas, 1994; Fujiwara *et al*., 2021), and so on.

It is known that activated area in a reaction-diffusion pattern is approximately proportional to total size even though it can be affected by boundary condition as discussed in (Gierer and Meinhardt, 1972) and reviewed in (Chapek and Müller, 2019). Therefore, actual positional information usually sets into the intended domain as an effect of scaling. Though, the positional information in this study was a uniaxial gradient that ignore the size of the object, the characteristic shape of additive growth became obvious when the boundaries grow over the lengths of the gradients. In Fig. S1, we show a case of shift of the positional information from without (*w_y_* = 0) inhibition when the shape within length β to a gradient when it over the length. This trial did not affect the results obtained. Such positional information is exist especially in leaf developments (Kazama et al., 2010, Tsukaya, 2013, Nakayama et al., 2014). That is, in actual leaf, it seemed that cell proliferation dosen’t biased in their early stage of formation within a boundary of cell cycle arrest front (AF).

However, the same gradient became a generator of various shapes in the case of expansive growth implemented as an algorithm (Fig. 3D, E). Differences in initial conditions or in propagation speeds will also change the shapes obtained. Even in a combination with the additive growth, the expansive growth kept the capacity to generate shape variation, accompanied by a loss of the regulation to the gradient (Fig. 3F-I). Therefore, the additive always governed by the gradient, though, the expansive which can escape it. However, these shapes still maintain robustness if the set of parameters is fixed. A same shape can be obtained repeatedly by an appropriate adjustment of the set of parameters.

Entire leaves with different proportions were reproduced by the combination of both modes with a difference in lengths of graded positional information (Fig. 4). The molecular mechanism for such difference was suggested in (Kierzkowski *et al*., 2019), then we treated “growth” of different modes expressed by simple algorithms. The different modes are considered caused by difference in progressions of tissue differentiation, in where the corresponding cells were located. In this case the positional information will be read-out to the boundary between cell proliferative phase and cell expansions after determination, known as AF on leaves in *A. thaliana* (Donnelly *et al*., 1999; Kazama, *et al*., 2010). Several molecules on the leaf blade are known to be related to the boundary determination of the plate meristem (Tsukaya, 2021). There are several examples that shows perturbations of such positional information change leaf proportions (Horiguchi *et al*., 2005; Kawade *et al*., 2010). Then differences in size and shape among other simple leaves are known to be derived from allometric growth patterns along with proximo-distal axis (Gupta & Nath, 2015). It was considered that the difference in proportions might be caused by the axially biased expansions at the different ranges, though, the combination of the boundary growths was not simple, i.e., a combination of a biased periodicity and graded uniaxial expansions.

As a biological relevance of this model, the hebetate protrusions that obtained by local growth in the additive growth algorithm (Nakamasu *et al*., 2014) can be observed in actual leaflets (Gleissberg, 2004, Ikeuchi *et al*., 2013) and in younger tooth in *A. thaliana* (Kawamura *et al*., 2010). *Papaveraceae* primordia with a certain difference in cell sizes shows such shapes in leaflets in early organogenesis stage (Gleissberg *et al*., 2004). Thought, it might not be derived from the difference in the tissue maturation states as mentioned in (Ikeuchi *et al*., 2013). Therefore, almost similar sizes of cells on the tips of the primordia might be caused by repeated cell proliferations that were described by the additive growth algorithm. Furthermore, the combined growth modes confirmed as mixed cell sizes is known to result in pointed protrusions observed in leaf serration in *A. thaliana* (Kawamura *et al*., 2010). Similar pointed shapes can be observed in unicellular algae, then it was treated with decrease in spacing in a developmental model (Laccalli and Harrison, 1987; Holloway and Harrison, 1999). However, the cases with bias had not been investigated. D’Arcy Thompson dealt with leaf shape in chapter IX of his book “On Growth and Form” (Thompson, 1917). In this chapter, he said that the balance between radial and tangential growth velocities is important for the leaf shapes exemplified in the Fig. 127. The present algorithm for biased expansion seems to affect this balance.

It was expected that the expansive growth in this study included distortions caused by the bias as shown in Fig. 8. Magnification of a triangle gives equal extensions of the three sides; however, it is obvious that magnified triangles with different expansion rates cannot connect their adjacent apical edges, even though they share basal apices at the geometric center. In closed contour, these apical edges need to be connected. As shown in Fig. 5A, polygons with different growth rates in its sides always have slants. i.e., an isosceles shown as meshed triangle in Fig.5A has slant. It can observe in serrations but it is uncertain whether it occur in formation of entire leaves. Though the whole shape in schematics of simulations (Fig.5A) has sharper distal and blunter proximal as frequently sown in leaf shapes. When we pick up arbitral one of divided triangles in a polygon (Fig. 5B), the apical edge propagates with some slants. The degree of slants will different with degree of the center angles, propagation speeds, and position of the triangle, dependently. As a results, generated differences in edge lengths and interior angles lead to slants in the shape. These problems yet have less biological relevance so they need to be addressed in more detail.

**Fig. 5.**
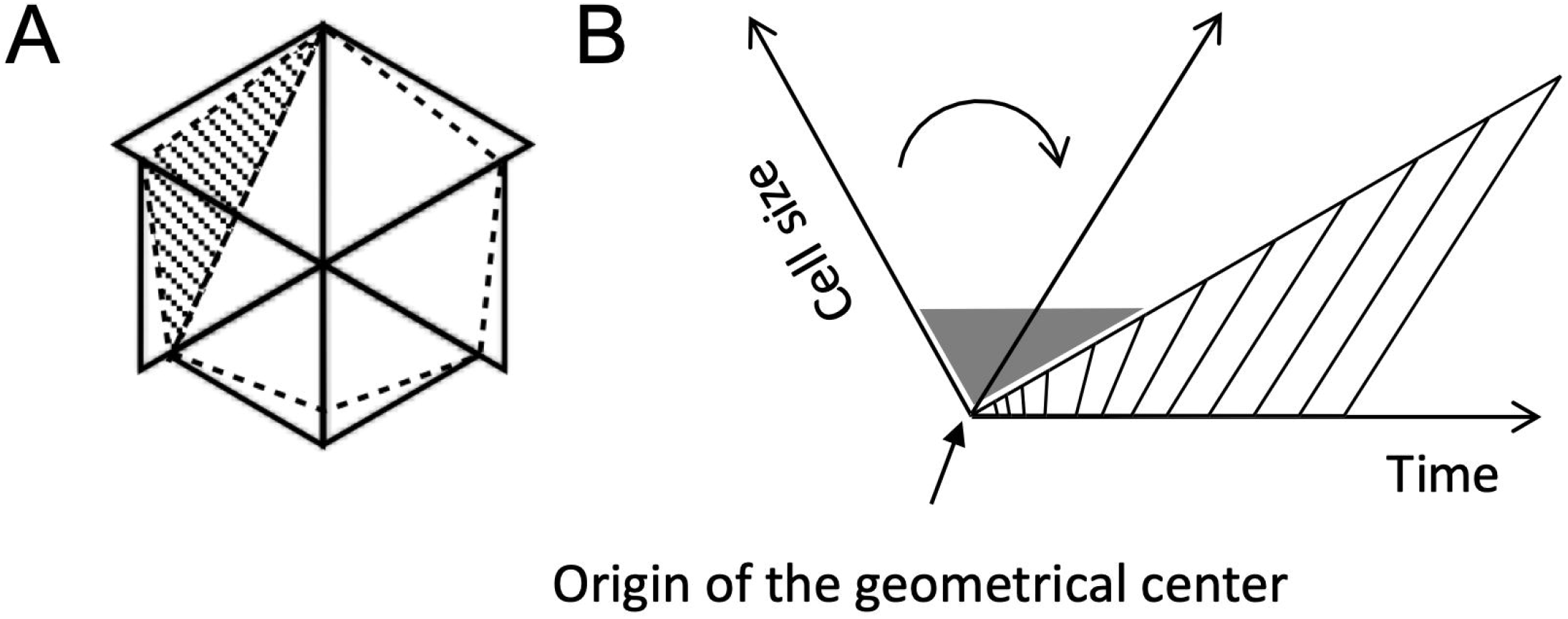
Illustration of slants included in biased expansive growth. (A)Superimposed magnified triangles with graded growth rates and expected results of biased expansion in this study. Former are shown by solid lines, and the latter are indicated by dashed lines. An initial isosceles (meshed triangle) become slant with change in lengths of the sides. (B) One of the divided triangles in a polygon. The apical edge propagates with some slants in the gradient (gray shade). The change in the directions of edges (the points at a start and when the edge escaped the gradient) were shown by two axes with a curved allow.

## Conclusion

Different algorithms for two types of growth modes brought different responses against simple positional information, i.e., that always governed by it or that can escape it. It was predicted theoretically that an expansive growth has a capacity to become a generator of oblong diversity in leaf morphogenesis. The effect was confirmed even in the combination case of additive growth with negate effects.

## Auther contribution

The author confirms being the sole contributor of this work and has approved it for publication.

## Acknowledgements

Author thanks Dr. T. Higaki, members of his laboratory, and IROAST staffs for the research environment provided. Thanks also to Dr. S. Kimura and Dr. N. J. Suematsu for the repeated discussions on leaf-shape morphogenesis, and Dr. A. Mochizuki for comments on initial conditions. Then I’d like to thank to anonymous reviewer(s) for kind reviews of my manuscript. Then I’d like to thank this beautiful nature with abundant diversity.

## Data availability statement

The raw data supporting the conclusion of this article will be made available by the authors, without undue reservation.

